# Predicting genomic traits in ammonia-oxidizing archaea using phylogenetic signals

**DOI:** 10.1101/2023.09.14.557535

**Authors:** Miguel A. Redondo, Christopher M. Jones, Pierre Legendre, Guillaume Guénard, Sara Hallin

## Abstract

Phylogenetic conservatism of microbial traits has paved the way for phylogeny-based predictions, allowing us to move from descriptive to predictive functional microbial ecology. Here, we applied phylogenetic eigenvector mapping, an approach not previously used for microorganisms, to predict key traits of ammonia-oxidizing archaea (AOA), which are important players in nitrogen cycling. Using 168 nearly complete AOA genomes and metagenome assembled genomes from public databases, we predicted the distribution of 18 ecologically relevant genes across an updated *amoA* gene phylogeny, including a novel variant of an ammonia transporter found in this study. Of the selected genes, 94% displayed a significant phylogenetic signal and gene presence was predicted with >88% accuracy, >88% sensitivity, and >80% specificity. The phylogenetic eigenvector approach performed equally well as ancestral state reconstruction of traits. We implemented the predictive models on an *amoA* sequencing dataset of AOA soil communities and show key ecological predictions, e.g., that AOA communities in nitrogen rich soils have capacity for ureolytic metabolism while those adapted to low pH soils have the high affinity ammonia transporter (*amt2*). Predicting genomic traits can shed light on the potential functions that microbes perform across earth biomes, further contributing to a better mechanistic understanding of their community assembly.

## Introduction

Genomic information is essential for trait-based approaches in microbial ecology, which can provide mechanistic explanations to ecological dynamics and ecosystem functions [1]. Since many microbial functions can be directly inferred from genomic content and microbial traits are phylogenetically conserved [2, 3], there is a framework for phylogeny-based trait predictions [4, 5] in which we can use massive amounts of environmental sequence data to predict the potential functions that microorganisms perform in the environment. In this way, we can impute functions to taxa whose genomes are not yet available and characterize ecologically relevant groups that represent a minor fraction in metagenomes. Phylogeny-based trait imputation have generally used either ancestral state reconstruction (ASR) by means of phylogenetic generalized least squares [6–8] or phylogenetic eigenvector maps (PEM) [9–11]. In contrast to ASR, PEMs offer the additional advantage of accommodating different modes of evolution, as well as the possibility of using phylogenetic signals together with abiotic factors when predicting traits or species distributions [12]. While PEMs have been used in a wide range of studies for macro-organisms, they have yet to be applied to microbial communities.

Among microorganisms, ammonia-oxidizing archaea (AOA) are an optimal group to evaluate the power of PEMs for predictions of genomic traits. First, they are key players in the nitrogen cycle and inhabit most ecosystems on earth [13, 14]. Given their ecological relevance, AOA genomes and metagenome-assembled genomes (MAGs) are increasingly available, thus expanding our knowledge of the functions of archaeal genes [15–17]. Second, there is a coherence between the organismal phylogeny and that of the *amoA* gene encoding the ammonia monooxygenase, which has long been used as a marker gene for AOA in environmental studies [18–20]. This has resulted in the availability of a global *amoA* phylogeny [21] that reflects environmental preferences of the organisms at different scales [22–24]. Whether or not gene content can be predicted using the *amoA* phylogeny, thus providing a basis for a more mechanistic understanding of AOA community assembly, remains uncertain.

The aim of this work was to predict genomic traits in AOA using the *amoA* phylogenetic signal. To reach this goal, we updated a recent *amoA* gene reference phylogeny of AOA [21] by adding 168 highly complete AOA genomes and MAGs available in public databases. The updated phylogeny was then used together with phylogenetic eigenvector mapping [11] to predict the presence of a set of genes (Table 1) belonging to four functional categories selected from a comparative genomics study [16], and a gene encoding a novel ammonia transporter found in the present study. We validated the predictions using hold-out validations and compared them to estimations based on ancestral state reconstruction. Finally, we implemented the predictive approach on AOA communities obtained from a field study [25], and linked the predicted AOA genes to soil properties by means of simultaneous analysis of environmental characteristics, species distributions, and species traits [26]. This study revealed that PEMs are useful for highly accurate predictions of genomic traits in AOA and can inform about their potential functions in the environment and the mechanisms underpinning community assembly.

**Table 1.**
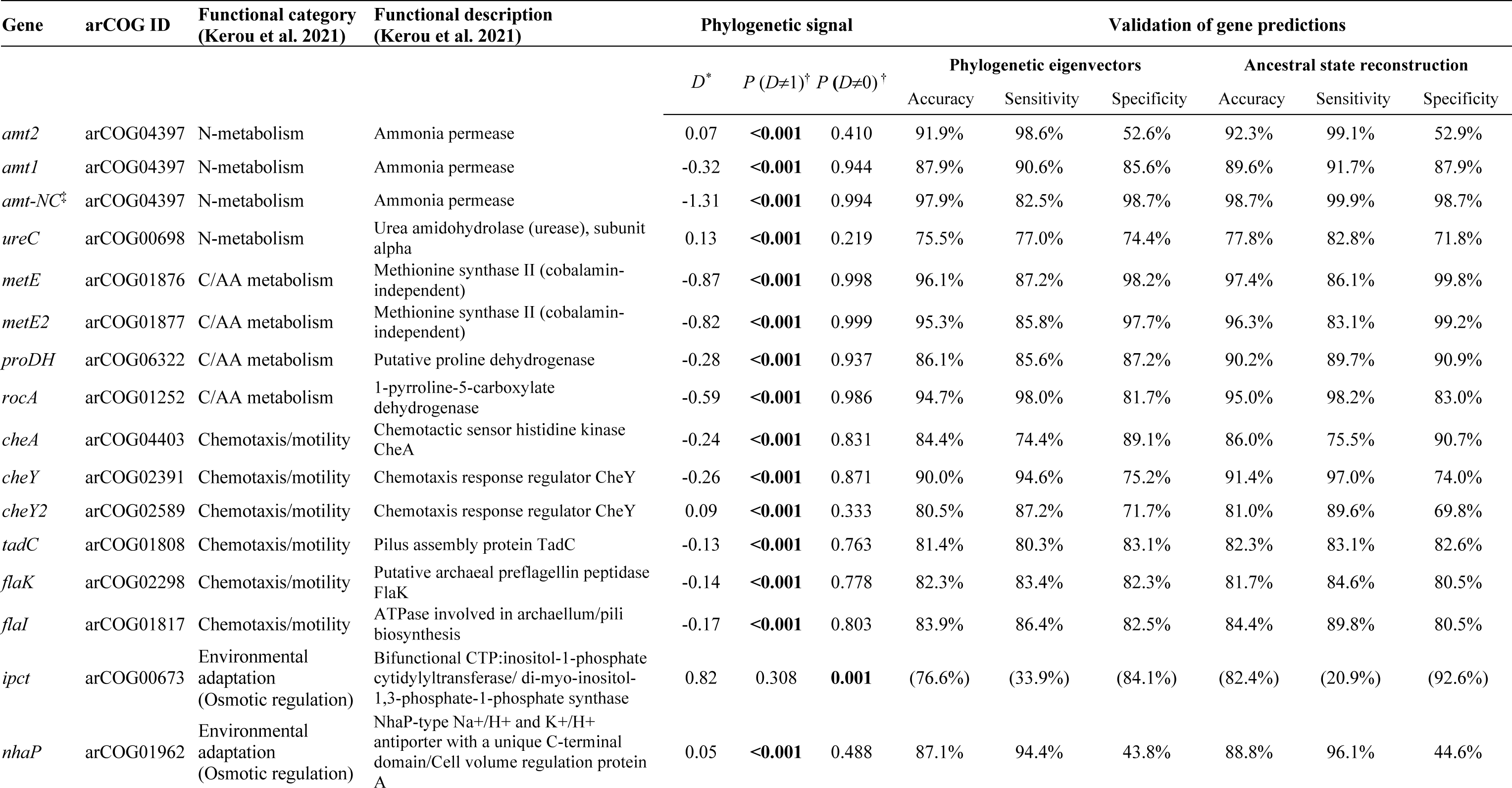
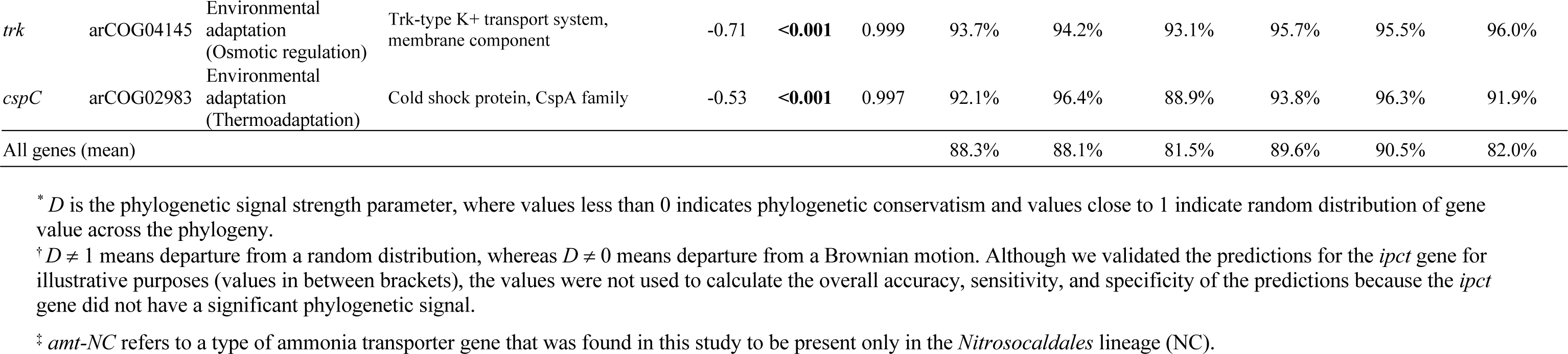
Genes selected for phylogenetic modelling, statistics for phylogenetic signal test, and validation results using phylogenetic eigenvectors and ancestral state reconstruction.

## Methods

### Update of the archaeal amoA reference phylogeny

We updated the *amoA* phylogeny developed by Alves et al. [21] by including all AOA genomes from isolates and metagenome-assembled genomes (MAGs) downloaded from the National Centre for Biotechnology Information (NCBI; https://www.ncbi.nlm.nih.gov/) and the Joint Genome Institute (JGI; https://genome.jgi.doe.gov/portal/), up to October 2021. Sequences of 1527 archaeal genomes and MAGs belonging to phylum Nitrososphaerota were initially screened for the presence of *amoA* genes using HMMER (http://hmmer.org) and the translated alignment from Alves et al. [21] as seed alignment. Significant hits (e-value < 10^-6^) were then aligned by amino acid to the original seed alignment using HMMER. An initial phylogeny of the significant hits, together with the 1190 sequences from Alves et al. [21] and the *amoA* sequences of seven ammonia-oxidizing bacteria to be used as an outgroup, was generated from the nucleotide alignment using FastTree 2.1 [27] to ensure the correct identification of AOA *amoA* genes. The alignment and tree were then inspected using the ARB software [28] to correct alignment errors and remove fragmented or poor quality (i.e. multiple ‘N’s) sequences, resulting in a total of 457 genomes and MAGs with *amoA* genes. We further discarded genomes/MAGs with <80% completeness or >5% contamination as determined by BUSCO [29] as well as those with identical *amoA* sequences and the same presence/absence values of the selected genes (see below) to remove duplicate taxa. At the end, *amoA* sequences from 168 high-quality genomes and MAGs were added to the original alignment of Alves et al. [21], and the total alignment was used to build maximum likelihood phylogenies using the IQTree software [30]. Tree search was carried out using ten independent tree searches, in which the first half used the default parameters while, for the second half, the ‘nbest’ and ‘nstop’ arguments were increased to 10 and 500, respectively. Node support for all trees was determined using ultrafast bootstrap approximation (1000 replicates). The ten trees were checked individually, and the tree with both the highest likelihood and coherence with the Alves et al. [21] topology was selected as the final reference tree. The model of the best tree was GTR+F+R10. The display of the phylogenetic trees for the present article was produced using iTOL [31].

### Selection of genes for predictions and screening of genomes

We selected 18 genes for phylogenetic modelling (Table 1) based on the comparative genomic study by Kerou et al. [16]. These genes were i) distributed across different AOA lineages to avoid genes only present in an isolated clade within the phylogeny, and ii) involved in pathways relating to different ecological functions and especially relevant to soil AOA. The selected genes belonged to four functional categories: nitrogen metabolism (*amt* and *ureC* genes), carbon and amino acid metabolism (*metE*, *proDH*, *rocA* genes), chemotaxis and motility (*cheA*, *cheY*, *flaK*, *flaI*, *tadC* genes) and environmental adaptation (*ipct, nhaP*, *trk* and *cspC* genes). We downloaded the orthologous gene protein alignment from each of these genes from the EggNOG 5.0 database (http://eggnog5.embl.de) using the arCOG ID reported by Kerou et al. [16]. When more than one arCOG was provided, we used each of them separately. With the alignments obtained from the EggNOG database, the 168 AOA genomes/MAGs were screened for the presence/absence of each gene using HMMER. To ensure that gene hits were accurate, we retrieved the protein sequences found on the genomes/MAGs, aligned them with the reference protein alignment from the EggNOG database and constructed phylogenies using the fastTree2 software [27]. The location of the hits relative to the reference sequences within the phylogenies was examined to make sure that they were in fact hits of the searched genes and not artefacts or other homologues with different functions. In the specific case of the ammonium transporter gene (*amt*), and due to its ecological relevance, the gene phylogeny was used to further classify the hits as Amt2 (high affinity) or Amt1 (low affinity) type of transporter [32–34]. The hits falling together with *Nitrosocosmicus* taxa in the phylogeny were assigned the low affinity transporter type (Amt1), as all sequenced *Nitrosocosmicu*s taxa encode uniquely one low affinity Amt [35–38]. It should be noted here that other studies have used the reverse nomenclature, in which Amt1 and Amt2 are defined as high and low affinity transporters, respectively [39–41]. To clarify these inconsistencies in the nomenclature, we screened the Amt hits of our study for the primers used by Nakagawa and Stahl [33] for both types of transporters, and found that the *amt2* primers from Nakagawa and Stahl [33] matched the genes that were defined as *amt1* in Offre et al. [39] and vice-versa. Therefore, we refer here to the gene encoding the high affinity transporter as *amt2* [32, 33], which corresponds to the *amt1* in Offre et al. [39]. The procedure of checking the *amt* hits using phylogenies allowed us to find a third type of ammonium transporter uniquely present in the *Nitrosocaldales* lineage (Supplementary Text 1; Supplementary Fig. 1-2; Supplementary Table 1). The gene names and arCOG references are provided in Table 1.

### Test of phylogenetic signal for binary traits

We tested the phylogenetic signal of each gene using the phylo.d function from the caper package [42], running 1000 permutations. This method determines the strength and the statistical significance of the phylogenetic signal for binary traits compared to random and Brownian motion distributions of the trait [43]. The resultant parameter (*D*) equals 1 when the binary trait has a random distribution across the phylogeny, and 0 if under Brownian motion. *D* values can be < 0 if the trait distribution is more clustered than expected under Brownian motion, and > 1 if it is more overdispersed than expected by random. The phylo.d function tests *D* for significant departure from 1 when the trait is phylogenetically conserved, and a significant departure from 0 when the trait does not follow a Brownian evolutionary model.

### Gene predictions using phylogenetic eigenvectors and comparison to ancestral state reconstruction

The input for the gene predictions was the final *amoA* phylogenetic tree and the matrix of presence/absence of the 18 genes across the 168 AOA genomes/MAGs. For gene predictions, we used phylogenetic eigenvectors obtained from the phylogenetic tree and compared the prediction results to those obtained by ancestral state reconstruction. In both cases, we predicted the presence of each gene in the taxa across the *amoA* phylogeny that had unknown genomes.

The phylogenetic eigenvector-based predictions were done in three steps: 1) decompose the phylogenetic tree into eigenvectors, 2) fit and regularize individual predictive models for each gene using the eigenvectors as the descriptors, 3) estimate the presence/absence of the genes from the models of step 2 on the taxa with unknown genomes given their locations in the phylogeny. To decompose the phylogenetic tree into eigenvectors (step 1), we used R package MPSEM [44]. The input for the MPSEM package is the phylogenetic tree containing all tips, i.e., taxa with and without gene information. The MPSEM package calculates an influence matrix using only the tips of the tree for which there is information about presence/absence of genes by pruning from the tree the taxa without trait information. From the influence matrix, phylogenetic eigenvectors were obtained by singular value decomposition. When calculating the phylogenetic eigenvectors, we fixed argument a=0 to assume a Brownian motion evolution of all traits. In step 2, the phylogenetic eigenvectors were then used as fixed factors in a multiple logistic regression whose coefficients were regularized using elastic net regularization, which combines *L_2_* (ridge regression) and *L*_1_ (LASSO) penalties on the model coefficients. For model regularization, we used the R package glmnet [45] with arguments family=“binomial” and α = 0.5 to use same amounts of both *L*_1_ and *L*2 shrinkage. The penalization hyper-parameter (*λ*) was tuned using leave-one-out cross validation within the training dataset and choosing the *λ* value that provided the highest accuracy of the predictions. The outcome of step 2 is a regularized model that predicts the probability of gene presence given the eigenvectors’ values. We classified the probabilities of the predictive model into presence or absence by choosing a threshold that maximizes both true positive (sensitivity) and true negative rate (specificity) of the predictions. For this, we created ROC curves using the R package pROC [46] with the function roc, and used the function coords to select a threshold for classification that would render the highest Youden’s J statistic, where J = sensitivity + specificity – 1. Probabilities that had values equal to or greater than the selected threshold were classified as presence, whereas probabilities lower than the threshold were classified as absence. The final output of step 2 is a model with tuned *λ* and classification threshold parameters that predict the gene presence for a taxon given its phylogenetic eigenvector scores. We then obtained the eigenvector scores for taxa with unknown genomes and used them on the model of step 2 to predict the gene presence. Function getGraphLocations found in MPSEM places the taxa with unknown genomes in the initial influence matrix of step 1 and function Locations2PEMscores obtains the phylogenetic eigenvector scores. It is important to note that the influence matrix and phylogenetic eigenvectors are not obtained using all tips of the tree, but only the ones with known gene information. For taxa with unknown information, we calculated the eigenvector score values, which are projections of new influence matrix coordinates on the eigenvectors obtained from the initial influence matrix (see Guénard et al. [11], for details on this procedure). Thus, the number of phylogenetic eigenvectors of the training dataset, and therefore the number of coefficients on the predictive model, is independent of the number of taxa to be predicted. The code can be used for analysis of other datasets by providing the phylogenetic tree and its associated binary trait table with and without missing values (NAs) in the input data.

The ancestral state reconstruction was done with R package picante R [47]. We used the phyEstimateDisc function to predict the genes of the AOA taxa with unknown genomes. In this procedure, for each taxon with unknown trait data, the phylogenetic tree is rerooted on the most recent ancestor common to the unobserved taxon and the rest of the phylogeny. The trait of the unobserved taxon is then estimated from the ancestral state reconstruction of the root of the rerooted phylogeny [6, 48]. The function phyEstimateDisc provides a trait state, i.e., presence or absence, for a given threshold (default = 0.5), as well as a value for the statistical support of the trait state.

### Validation of the predictive models

We validated the predictions of each gene using a 20% hold-out validation. This procedure consists of randomly removing 20% of the initial dataset before creating the predictive model and validating it on the removed samples. We repeated this procedure 500 times, always randomizing the taxa included in the held-out dataset. For each gene, we obtained the mean accuracy, sensitivity, and specificity of the prediction. We also varied the proportion of taxa to be held out in the validations to 30% and 40%, and obtained accuracies of 87.1% and 86.4%, respectively (compared to >88% accuracy at 20% hold out). We could not increase the proportion of data to be held out because many genes were class unbalanced. Both sensitivity and specificity were obtained using R package pROC.

We validated the accuracy of the predictions at the genome/MAG level using leave-one-out cross validation, in which we deleted each genome/MAG from the dataset in turn and used the rest of the genomes to predict the gene content of the previously removed genome or MAG.

### Implementation of predictive modelling on natural communities: a case study

To illustrate how predicting AOA traits could help scientists link community composition, predicted traits, and environmental properties, we used data from a previous study that characterized AOA communities by *amoA* amplicon sequencing on 50 sampling points across an agricultural area in which soil properties were measured (see Enwall et al. [49] and Jones & Hallin [25] for more information). The study site is a 44 hectare farm divided into 14 fields, with sampling points taken at 51 locations throughout the field based on environmental gradients identified in a previous study [50]. We deleted one of the locations (S17) because it lacked data on soil properties. We predicted the presence/absence of the 17 genes showing significant phylogenetic signal on the AOA communities and linked the gene composition with the soil properties that were reported by Enwall et al. [49].

To predict the traits of the AOA members of the soil communities, we placed the *amoA* sequences of each OTU (162 in total) on the reference phylogeny using the EPA-ng [51] (accessed at https://github.com/pierrebarbera/epa-ng) and gappa software [52] (accessed at https://github.com/lczech/gappa), therefore obtaining a new phylogenetic tree containing all sequences of the updated reference phylogeny and the representative OTU sequences to be predicted as pendant branches. We used this phylogenetic tree to implement the phylogenetic eigenvector modelling described above. The output was a matrix with presence/absence of all 17 genes for each OTU in the study.

We studied the link between the predicted genes and the soil properties mediated by community composition by performing a RLQ analysis followed by a univariate fourth-corner analysis [26, 53, 54]. RLQ is an ordination method that displays the covariation of traits, i.e., the predicted genes, and environmental properties providing site and species scores, and a global test for significance. The fourth-corner correlations, on the other hand, provide tests of single associations between genes and environmental properties. The two methods are complementary and can be performed sequentially. Following Dray et al. [26], we calculated three separate ordinations for the RLQ analysis: a correspondence analysis for the community data, a principal component analysis after standardization of the variables in the environmental data matrix, and a principal component analysis for the predicted genes presence/absence without standardization. The three ordinations were analyzed together using the rlq function of the ade4 package for R [55]. The RLQ analysis was tested for significance using the randtest function of the ade4 package with argument modeltype = 6 for the permutation test. This model performs two sequential permutational tests, a first one testing the link between species distribution and environmental conditions (model 2), and a second testing the link between species distribution and traits (model 4). When both tests are significant, the highest of the two *P*-value provides the statistical significance for the global tests of association between traits and environment [56].

To test which specific gene was associated with each soil property, we performed a fourth-corner analysis using the fourthcorner function from the ade4 package, with modeltype = 6, 99 999 permutations, and “false discovery rate” as the multiple testing correction method for the *P*-value.

## Results

### Congruence between amoA phylogeny and content of genes for predictions

To perform the phylogenetic eigenvectors-based predictions, we first updated the archaeal *amoA* gene reference phylogeny from Alves et al. [21], which now contains 1190 unique *amoA* sequences and 168 highly complete AOA genomes of isolates and MAGs distributed across most *amoA* lineages (Supplementary Fig. 3). We screened the genomes and MAGs for the presence/absence of a set of ecologically relevant genes selected for modelling [16] (Table 1). We found a negative association between *Nitrososphaerales* (NS) taxa and the presence of genes associated with motility (*tadC*, with the exception of *NS*-α), osmotic regulation (*trk*), and thermoadaptation (*cpsC*). The *metE* gene, responsible for methionine synthesis in energy-limiting environments [16] was found in *Nitrosopumilales* (NP) clades associated to deep sea waters (*NP*-α and -θ), as well as in *Nitrosocaldales* (NC) and Nitrosotaleales (NT). Genes encoding the high-and low-affinity ammonium transporters (Amt2 and Amt1, respectively [32, 33], see details in Methods) were identified based on their positions within a phylogenetic tree of translated Amt sequences [39]. When doing this, we identified a gene encoding a novel variant of the ammonia transporter protein specifically associated with the *NC amoA* lineage, hereafter referred to as Amt-NC. All taxa having the *amt-NC* gene, also had the *amt*2 (Fig. 1a). We found two subgroups of the ammonium transporter gene *amt-NC*, hereinafter *amt-NC.1* and *amt-NC.2* (Supplementary Fig. 1), which corresponded to the groups into which Luo et al. [17] divided the *NC* lineage based on the concatenation of 122 archaeal genes (Supplementary Text 1). The amino acid composition of the Amt-NC was identical to the Amt types described by Offre et al. [39] at the ammonium-binding sites, i.e. they contained the same histidine lining the transporter pore [39] and had the same amino acids in several conserved loci (Supplementary Fig. 2). However, it differed in several loci across the regions described by Offre et al. [39] (Supplementary Fig. 2).

**Figure 1.**
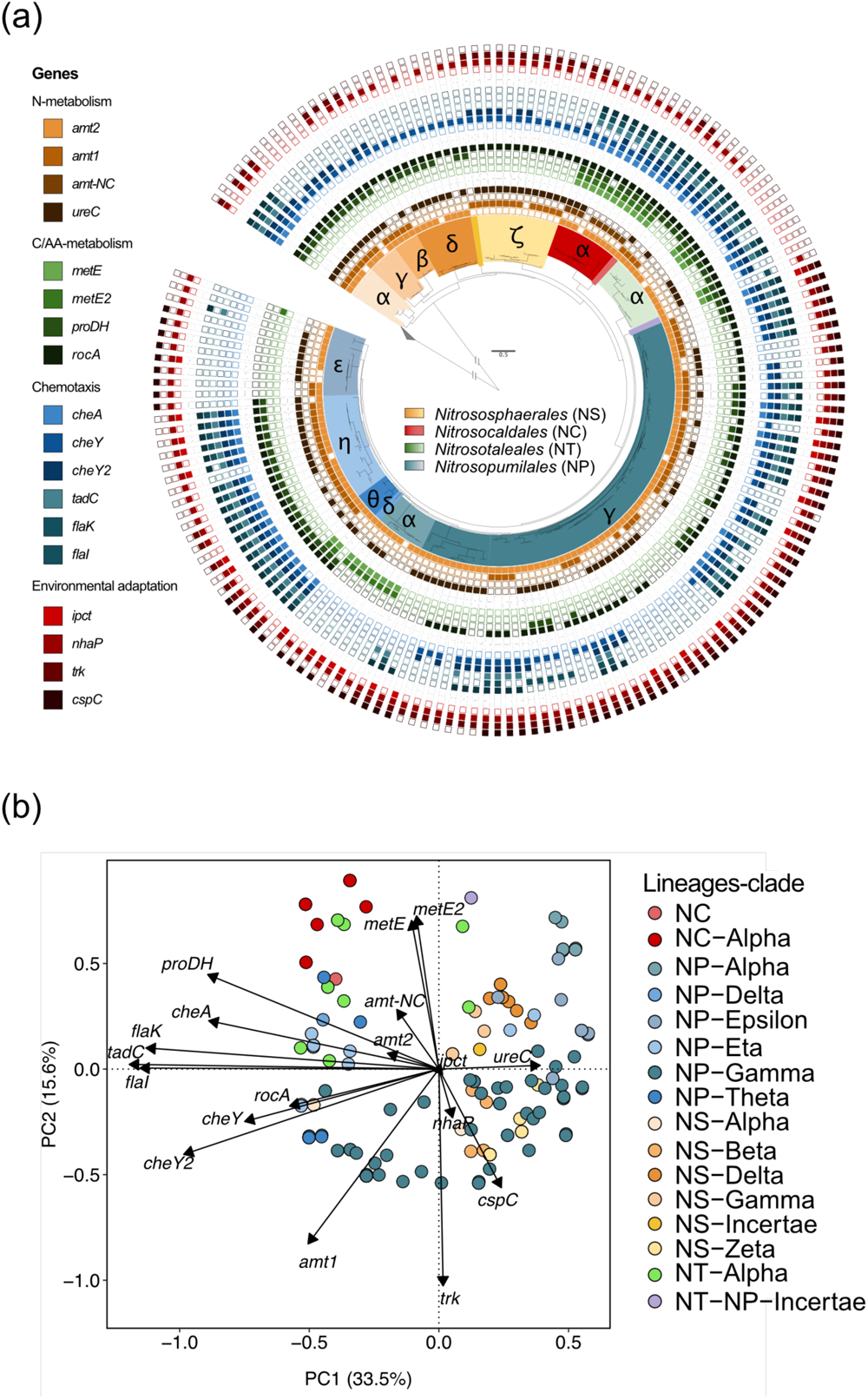
Distribution of genomic traits in AOA genomes. (**a**) The presence/absence of 18 ecologically relevant genes (Table 1), across a phylogeny of 168 AOA isolate genomes and MAGs. Colors in outer rings denote presence (filled squares) and absence (empty squares) of specific genes. Clades within each AOA genus are denoted by Greek letters and the tree is rooted on a collapsed gray clade of bacterial *amoA*. (**b**) Principal component analysis of the 168 genomes and MAGs based on a presence-absence matrix for the 18 selected genes.

All selected genes except *ipct* were phylogenetically conserved and did not depart from a Brownian motion model (Table 1). The distance between the 168 isolate genomes and MAGs in a principal component ordination including all 18 genes reflected the lineage classification of the AOA taxa (Fig. 1b), supporting a coherence between the *amoA* phylogeny and the overall gene content of available genomes and MAGs.

### Phylogeny-based predictions of selected genes

We used the gene content and phylogenetic relatedness of the 168 genomes and MAGs to build predictive models of gene presence. For each of the 18 genes, we used elastic net regularized regressions, with the PEMs as predictors and the presence of each gene as response. The models with optimized lambda penalization and classification threshold parameters (Supplementary Table 2) were then used to predict gene presence across the *amoA* phylogeny using the phylogenetic eigenvector scores of unobserved taxa as input. The predicted presence of most genes varied across and within *amoA* lineages. For example, genes encoding the high-affinity ammonia transporter (*amt2*) was predicted to be present in nearly all lineages except two *NS* groups, *NS-*ρ. and NS-σ. In contrast, the low affinity ammonia transporter (*amt*1) and urease (*ureC*) genes were predicted to be within most *NS* clades, yet were more unevenly distributed or absent across *NP* clades (Fig. 2). Motility genes showed the highest within-clade variation, where presence of genes related to archaellum formation (*flaK* and *flaI*) varied between taxa of the *NP*-γ clade (Fig. 2), while nearly all lineages except *NP*-ρ, and -α were predicted to have the *cheY* gene, encoding a response regulator associated with chemotaxis. The gene responsible for B-12 independent methionine synthesis (*metE*) was predicted to be present in all *NC* and in most *NT* taxa. Across *NP* clades, *metE* was predicted to be present in NP-α and -β, as well as in most *NP*-θ and some *NP*-ρι, -ρ,, and -ψ (Fig. 2). Genes associated with osmotic regulation (*nhaP*, *trk*) and thermoadaptation (*cpsC*) were also predicted to be present more often in *NP* clades and less often in *NS* (Fig. 2).

**Figure 2.**
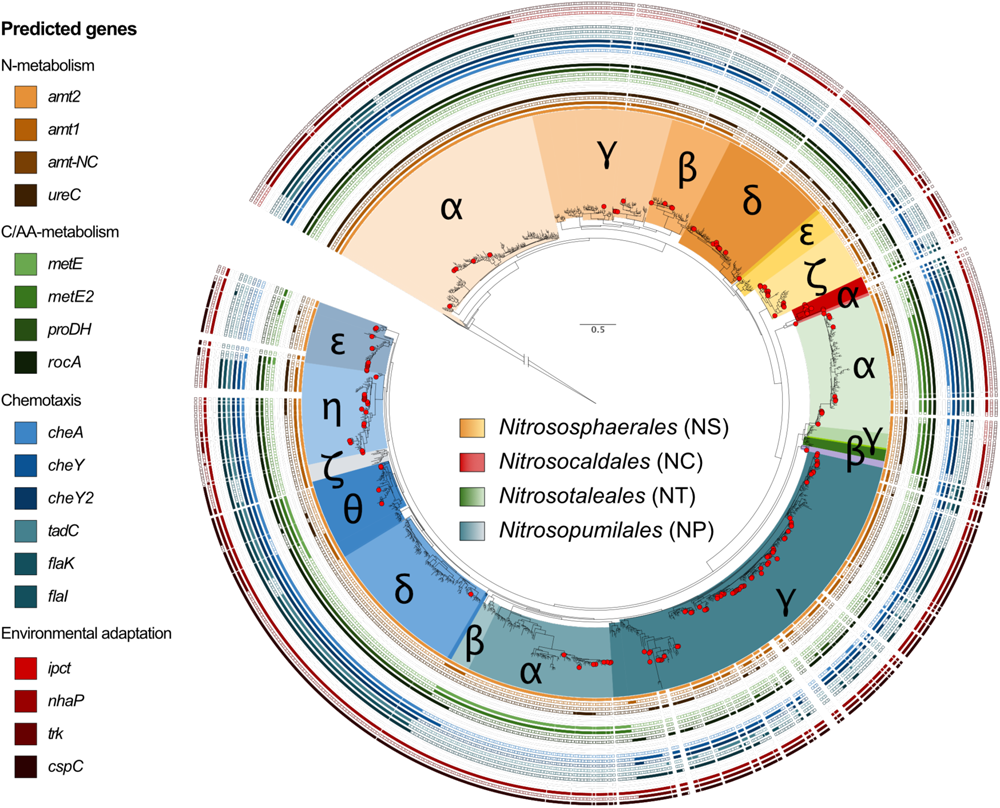
Predictions of gene presence/absence across the archaeal *amoA* reference phylogeny. Colors in outer rings denote predicted presence (filled squares) and absence (empty squares) of specific genes (see Table 1 for definition). Red circles are the isolate genomes and MAGs with > 80% completeness and < 5% contamination. Clades within each AOA genus are denoted by Greek letters and the tree is rooted on a collapsed gray clade of bacterial *amoA*.

The PEM-based predictions of gene presence for all genes resulted in an average of 88.3% accuracy, 88.1% sensitivity, and 81.5% specificity based on a 20% hold-out-validation (Table 1). Ancestral state reconstruction of traits resulted in similar levels of accuracy, sensitivity and specificity of predictions (89.6%, 90.5%, and 82.0% respectively; Table 1). For both methods, the predictive accuracy increased linearly with the strength of the phylogenetic signal (R^2^ = 0.64 for both the PEM and ancestral state reconstructions, Supplementary Fig. 4).

### Link between predicted genes and soil properties; a case study

To exemplify how gene predictions can contribute to trait-based studies, we placed the *amoA* sequence of the members of 50 AOA communities from arable land in the updated *amoA* reference phylogeny, predicted the gene presence among these *amoA*-based OTUs (Fig. 3a), and tested the link between predicted genes and soil properties by performing RLQ [53] and fourth-corner [54] analyses. The first RLQ axis showed that OTUs predicted to have genes encoding the high-affinity ammonia transporter (*amt*2) and chemotaxis response (*cheY*2) and lacking the *nhaP* gene, were more associated with sites with lower pH (pH = 5.7 - 6.0 in sites S19, 21, 23, and 28; Fig. 3 b, c). The second RLQ axis highlighted two specific genes, i.e., the low-affinity ammonia transporter (*amt*1) and the *ureC*, which were most closely associated to sites with higher levels of total nitrogen and carbon, higher ammonium content, and more moderate soil pH (pH = 6.1 - 6.2 in sites S29, 31, 34, 37; Fig. 3b, c). When testing the univariate associations of each gene and soil property using the fourth-corner approach, we found a negative correlation between the potential ammonia oxidation activity in soils and the presence of the high-affinity ammonia transporter (*P* = 0.04, after false discovery rate correction; Supplementary Fig. 5), and a positive correlation between the presence of the *ureC* gene and total nitrogen (*P* = 0.06, after false discovery rate correction; Supplementary Fig. 5).

**Figure 3.**
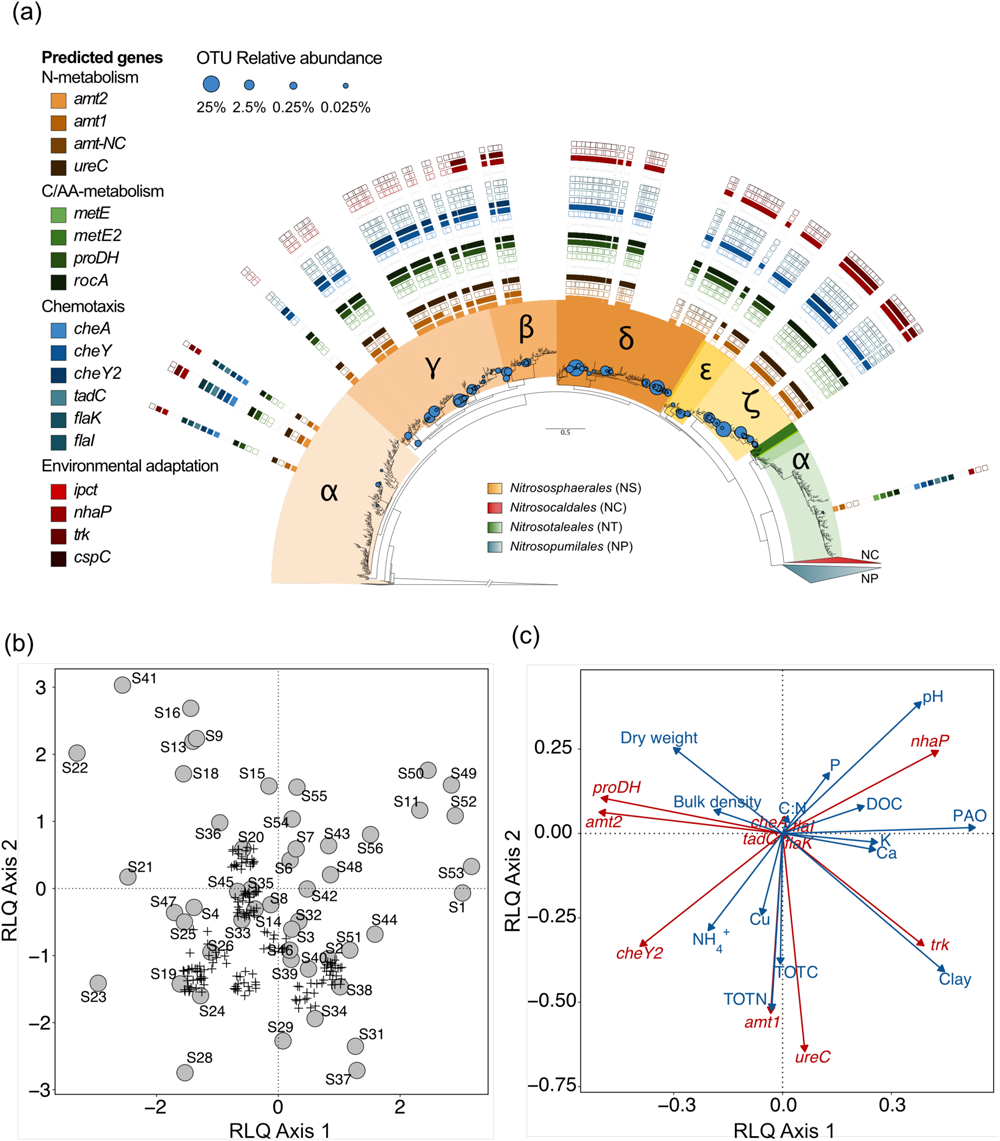
Association of predicted genes of AOA communities with soil properties across 44 ha of arable land. (**a**) Placements of the 162 Operational Taxonomical Units (OTUs) on the *amoA* reference phylogeny. Each circle represents an OTU whose size is proportional (fifth root) to their relative abundance. Colors in outer rings denote predicted presence (filled squares) and absence (empty squares) of specific genes for each OTU (see Table 1 for definition). None of the OTUs belonged to *Nitrosopumilales* nor *Nitrosocaldales*, and therefore both lineages are collapsed in the tree. The tree is rooted on a collapsed gray clade of bacterial *amoA*. **(b)** Biplot of the RLQ analysis displaying scores of 50 sites (S1-S50) and 162 Operational Taxonomical Units (OTUs) in circles and plus (+) signs, respectively. OTU symbols were jittered for visualization purposes. (**c**) Biplots of the RLQ analysis displaying association between predicted genes (red) and observed soil properties (blue). PAO refers to potential ammonia oxidation rates. The genes whose predicted values were all 0 or 1 were not included in the RLQ analysis. The global *P*-value associated with the RLQ analysis was 0.009.

## Discussion

Predicting genomic traits can inform about key functions that microorganisms perform in the environment without access to their full genomes. In this study, we predicted genes of ammonia-oxidizing archaea (AOA) with >88% accuracy using the *amoA* phylogenetic signals implemented in phylogenetic eigenvectors. The overall distribution of our isolate genomes and MAGs across the phylogeny makes us confident that these predictions are reliable, particularly for AOA belonging to clades with good genome representation (see Supplementary Table 3). While genome and MAG incompleteness may limit inferences of microbial functions [57], we did not find any association between predictive accuracy and genome completeness (Supplementary Fig. 6a). Only a weak and non-significant trend suggesting that presence of genes in highly complete genomes were predicted with lower sensitivity and higher specificity than less complete genomes was observed (Supplementary Fig. 6b,c). Thus, with more incomplete genomes in the training datasets, predictions of gene presence may be more reliable than predictions of gene absence.

The high prediction accuracy can be explained by the strength of the phylogenetic signal of the screened genes. More than 90% of the screened genomic traits were conserved phylogenetically, in line with previous studies [16, 19], and the strength of the phylogenetic signal was positively correlated with the accuracy of the predictions [5, 8]. As an example, the *amt-NC* and *ureC* gene, which had the strongest and weakest phylogenetic signal, respectively, displayed the highest and lowest accuracy of predictions, respectively. The differences in phylogenetic signal between genes is likely to be the evolutionary result of niche specialization. The *amt-NC* gene, found here for the first time, encodes a variant of ammonia transporter present only in *Nitrosocaldus* AOA inhabiting thermal waters (Supplementary Text 1), and is therefore localized in a single clade within the *amoA* phylogeny. By contrast, the *ureC* gene tends to be phylogenetically dispersed. In agreement, soil AOA thrive in conditions in which ammonia is supplied slowly through mineralization of organic matter [58], and urease genes are abundant in Thaumarchaeota communities in oligotrophic marine environments [59]. The high accuracy of the predictions may also be the result of the availability of AOA genomes and MAGs across the *amoA* phylogeny. Accordingly, predictions on genomes and MAGs belonging to the *NS*-ζ and *NC*-α, *NP*-ρ,, and *NP*-α clades had the highest accuracies, and these AOA lineages were well-represented by genomes with close phylogenetic relatedness to the within-clade available relatives [5, 60] (Supplementary Table 3). These findings support previous claims that phylogenetic modelling of microbial traits should be restricted to groups of organisms with good genomic representation in the databases [4, 61].

Genomic trait predictions can provide a mechanistic understanding of community assembly. By implementing the gene predictions in multivariate models of AOA communities in soils with varying physical and chemical properties, we show that AOA communities adapted to low pH soils are more likely to have the high-affinity ammonia transporter (*amt2*) and chemotaxis gene *cheY2*. This makes sense given the low availability of ammonia at low pH, and is in line with previous studies showing down-regulation of bacterial chemotaxis genes at high pH [62]. By contrast, high pH sites were associated with OTUs predicted to have the *nhap* gene encoding a protein related to Na^+^/H^+^ antiporters, which are more important for homeostasis under neutral or alkaline conditions [63]. Resource rich sites with high amounts of total nitrogen and ammonium as well as soil organic carbon, located in areas with higher organic N addition [22], were related to the dominance of AOA with predicted potential for ureolytic metabolism (*ureC*). Nitrogen addition may increase the abundance of *ureC* genes [64], which in turn is associated with high amounts of ammonium as a result of urea hydrolysis [65]. Higher potential ammonia oxidation activity was linked to taxa with the low affinity ammonium transporter, likely because the high ammonium concentrations used in the activity assay favored the ammonia oxidizers with low ammonia affinity or those that prefer inorganic to organic N sources, including ammonia-oxidizing bacteria [66]. These patterns of AOA community assembly can have key implication for ecosystem functions, such as CO_2_ emission rates, as a result of urea hydrolysis [67]. Overall, our predictions on soil AOA communities show that the combination of genomic trait predictions with RLQ and fourth-corner analyses can shed light on mechanisms of community assembly and this approach can be expanded to other microbial groups than AOA to decipher ecological dynamics.

Phylogenetic eigenvector mapping can complement the broadly used ASR when performing phylogeny-based modelling. We show that both methods have similar values of accuracy, sensitivity and specificity of the predictions, and accuracy values between these two methods across all genes were highly correlated (*R^2^* = 0.97). Based on co-inertia testing, both predictions across the phylogeny were associated with a high *RV* coefficient value, i.e., a correlation metric between two multivariate sets of variables [10], of 0.81 (data non shown). When setting the threshold for classifying probabilities at 0.5 in our PEM models, which is the default for ASR in the R package picante [47], the *RV* increased to 0.89. The use of either PEM or ASR depends on the role of phylogeny in the analyses [68, 69]. When estimating microbial traits using only phylogenetic signals, both methods give similar results, with ASR not requiring the extra step of selecting or regularizing a model [8]. On the other hand, PEM can be used when modelling phylogenetic signal together with other factors, e.g. abiotic variables or gene co-occurrences, which is useful when estimating traits on partially incomplete databases [70, 71]. In addition, the procedure used by the MPSEM R [44]package to calculate phylogenetic eigenvectors can use phylogenetic networks as input. Thus, PEM can potentially be used for microorganisms for which horizontal gene transfer occurs, as well as for organisms that undergo reticulate speciation or hybridization [72]. Finally, the sets of latent descriptors produced by PEM can be used with other modelling approaches, such as support vector machines, gradient boost machines or artificial neural networks. Although some studies report a better performance of ASR over PEMs [73], perhaps because of model selection issues, the present study shows that both can provide similar accuracies, while PEM constitutes a more versatile tool for phylogenetic modelling.

In this study, we show that we can move towards genomic trait prediction in microbial ecology. Whereas the absence of many microbial genomes can limit the implementation of trait-based studies in microbial ecology, many microbial genes are conserved phylogenetically, and their presence can be predicted using such tools as phylogenetic eigenvectors and ancestral state reconstruction. Predictive modelling of microbial functions can provide useful information to understand how evolution shapes the genetic content of microorganisms, how that determines their distribution in the environment, and how that ultimately impacts ecosystem functions.

## Acknowledgements

This research was supported by the The Swedish Research Council (grant 2020-00434 to M.A.R.). We are grateful to Gabriel Dansereau of Université de Montréal for insightful discussions during the analysis of the data.

## Data availability

The datasets with gene content and code to replicate the predictive analyses will become publicly available on Github (https://github.com/RedondoMA/AOA_gene_predictions) upon publication.

## References

1. Lajoie, G. & Kembel, S. W. Making the most of trait-based approaches for microbial ecology. Trends Microbiol. 27, 814–823 (2019).

2. Martiny AC, Treseder K, Pusch G. Phylogenetic conservatism of functional traits in microorganisms. ISME J 2013; 7: 830–838.

3. Morrissey EM, Mau RL, Hayer M, Liu X-JA, Schwartz E, Dijkstra P, et al. Evolutionary history constrains microbial traits across environmental variation. Nat Ecol Evol 2019; 3: 1064.

4. Langille MGI, Zaneveld J, Caporaso JG, McDonald D, Knights D, Reyes JA, et al. Predictive functional profiling of microbial communities using 16S rRNA marker gene sequences. Nat Biotechnol 2013; 31: 814–821.

5. Goberna M, Verdú M. Predicting microbial traits with phylogenies. ISME J 2016; 10: 959–967.

6. Garland, T. & Ives, A. R. Using the past to predict the present: Confidence intervals for regression equations in phylogenetic comparative methods. Am. Nat. 155, 346–364 (2000).

7. Kembel SW, Cowan PD, Helmus MR, Cornwell WK, Morlon H, Ackerly DD, et al. Picante: R tools for integrating phylogenies and ecology. Bioinformatics 2010; 26: 1463–1464.

8. Walkup J, Dang C, Mau RL, Hayer M, Schwartz E, Stone BW, et al. The predictive power of phylogeny on growth rates in soil bacterial communities. ISME Commun 2023; 3: 1–8.

9. Diniz-Filho JAF, de Sant’Ana CER, Bini LM. An eigenvector method for estimating phylogenetic inertia. Evol Int J Org Evol 1998; 52: 1247–1262.

10. Legendre P, Legendre L. Numerical Ecology. 2012. Elsevier.

11. Guénard G, Legendre P, Peres-Neto P. Phylogenetic eigenvector maps: a framework to model and predict species traits. Methods Ecol Evol 2013; 4: 1120–1131.

12. Guénard G, Lanthier G, Harvey-Lavoie S, Macnaughton CJ, Senay C, Lapointe M, et al. Modelling habitat distributions for multiple species using phylogenetics. Ecography 2017; 40: 1088–1097.

13. Stahl, D. A. & de la Torre, J. R. Physiology and diversity of ammonia-oxidizing archaea. Annu. Rev. Microbiol. 66, 83–101 (2012).

14. Lehtovirta-Morley LE. Ammonia oxidation: Ecology, physiology, biochemistry and why they must all come together. FEMS Microbiol Lett 2018; 365.

15. Kerou, M. et al. Proteomics and comparative genomics of *Nitrososphaera viennensis* reveal the core genome and adaptations of archaeal ammonia oxidizers. Proc. Natl. Acad. Sci. U. S. A. 113, E7937–E7946 (2016).

16. Kerou M, Ponce-Toledo RI, Zhao R, Abby SS, Hirai M, Nomaki H, et al. Genomes of Thaumarchaeota from deep sea sediments reveal specific adaptations of three independently evolved lineages. ISME J 2021; 15: 2792–2808.

17. Luo, Z.-H. et al. Genomic insights of “Candidatus Nitrosocaldaceae” based on nine new metagenome-assembled genomes, including “Candidatus Nitrosothermus” gen nov. and two new species of “Candidatus Nitrosocaldus”. Front. Microbiol. 11, (2021).

18. Leininger S, Urich T, Schloter M, Schwark L, Qi J, Nicol GW, et al. Archaea predominate among ammonia-oxidizing prokaryotes in soils. Nature 2006; 442: 806– 809.

19. Oton EV, Quince C, Nicol GW, Prosser JI, Gubry-Rangin C. Phylogenetic congruence and ecological coherence in terrestrial Thaumarchaeota. ISME J 2016; 10: 85–96.

20. Wang, H. et al. Linking 16S rRNA gene classification to amoA gene taxonomy reveals environmental distribution of ammonia-oxidizing archaeal clades in peatland soils. mSystems 6, (2021).

21. Alves RJE, Minh BQ, Urich T, Haeseler A von, Schleper C. Unifying the global phylogeny and environmental distribution of ammonia-oxidising archaea based on amoA genes. Nat Commun 2018; 9: 1–17.

22. Wessén E, Söderström M, Stenberg M, Bru D, Hellman M, Welsh A, et al. Spatial distribution of ammonia-oxidizing bacteria and archaea across a 44-hectare farm related to ecosystem functioning. ISME J 2011; 5: 1213–1225.

23. Gubry-Rangin C, Hai B, Quince C, Engel M, Thomson BC, James P, et al. Niche specialization of terrestrial archaeal ammonia oxidizers. Proc Natl Acad Sci 2011; 108: 21206–21211.

24. Dai S, Liu Q, Zhao J, Zhang J. Ecological niche differentiation of ammonia-oxidising archaea and bacteria in acidic soils due to land use change. Soil Res 2018; 56: 71–79.

25. Jones CM, Hallin S. Geospatial variation in co-occurrence networks of nitrifying microbial guilds. Mol Ecol 2019; 28: 293–306.

26. Dray S, Choler P, Dolédec S, Peres-Neto PR, Thuiller W, Pavoine S, et al. Combining the fourth-corner and the RLQ methods for assessing trait responses to environmental variation. Ecology 2014; 95: 14–21.

55. Price, M. N., Dehal, P. S. & Arkin, A. P. FastTree 2 – Approximately maximum-likelihood trees for large alignments. PLOS ONE 5, e9490 (2010).

28. Ludwig W, Strunk O, Westram R, Richter L, Meier H, Yadhukumar, et al. ARB: a software environment for sequence data. Nucleic Acids Res 2004; 32: 1363–1371.

29. Simão FA, Waterhouse RM, Ioannidis P, Kriventseva EV, Zdobnov EM. BUSCO: assessing genome assembly and annotation completeness with single-copy orthologs. Bioinformatics 2015; 31: 3210–3212.

53. Minh, B. Q. et al. IQ-TREE 2: New models and efficient methods for phylogenetic inference in the genomic era. Mol. Biol. Evol. 37, 1530–1534 (2020).

31. Letunic I, Bork P. Interactive Tree Of Life (iTOL) v5: an online tool for phylogenetic tree display and annotation. Nucleic Acids Res 2021; 49: W293–W296.

32. Wright CL, Lehtovirta-Morley LE. Nitrification and beyond: metabolic versatility of ammonia oxidising archaea. ISME J 2023; 17: 1358–1368.

28. Nakagawa, T. & Stahl, D. A. Transcriptional response of the archaeal ammonia oxidizer *Nitrosopumilus maritimus* to low and environmentally relevant ammonia concentrations. Appl. Environ. Microbiol. 79, 6911–6916 (2013).

34. Qin W, Zheng Y, Zhao F, Wang Y, Urakawa H, Martens-Habbena W, et al. Alternative strategies of nutrient acquisition and energy conservation map to the biogeography of marine ammonia-oxidizing archaea. ISME J 2020; 14: 2595–2609.

57. Alves, R. J. E. et al. Ammonia oxidation by the arctic terrestrial Taumarchaeote Candidatus *Nitrosocosmicus arcticus* is stimulated by increasing temperatures. Front. Microbiol. 10, (2019).

58. Lehtovirta-Morley, L. E. et al. Isolation of ‘Candidatus *Nitrosocosmicus franklandus*’, a novel ureolytic soil archaeal ammonia oxidiser with tolerance to high ammonia concentration. FEMS Microbiol. Ecol. 92, fiw057 (2016).

59. Sauder, L. A. et al. Cultivation and characterization of Candidatus *Nitrosocosmicus exaquar*e, an ammonia-oxidizing archaeon from a municipal wastewater treatment system. ISME J. 11, 1142–1157 (2017).

38. Jung M-Y, Kim J-G, Sinninghe Damsté JS, Rijpstra WIC, Madsen EL, Kim S-J, et al. A hydrophobic ammonia-oxidizing archaeon of the Nitrosocosmicus clade isolated from coal tar-contaminated sediment. Environ Microbiol Rep 2016; 8: 983–992.

39. Offre P, Kerou M, Spang A, Schleper C. Variability of the transporter gene complement in ammonia-oxidizing archaea. Trends Microbiol 2014; 22: 665–675.

61. Liu, L., Liu, M., Jiang, Y., Lin, W. & Luo, J. Production and excretion of polyamines to tolerate high ammonia, a case study on soil ammonia-oxidizing archaeon “Candidatus *Nitrosocosmicus agrestis*”. mSystems 6, 10.1128/msystems.01003-20 (2021).

62. Zou, D. et al. Genomic characteristics of a novel species of ammonia-oxidizing archaea from the Jiulong River Estuary. Appl. Environ. Microbiol. 86, e00736–20 (2020).

63. Orme, D. et al. caper: Comparative analyses of phylogenetics and evolution in R. (2018).

64. Fritz, S. A. & Purvis, A. Selectivity in mammalian extinction risk and threat types: a new measure of phylogenetic signal strength in binary traits. Conserv. Biol. 24, 1042–1051 (2010).

48. Guenard, G. & Legendre, P. MPSEM: Modeling phylogenetic Ssgnals using eigenvector maps. (2022).

65. Friedman, J., et al. glmnet: Lasso and elastic-net regularized generalized linear models. (2023).

66. Robin, X., et al. pROC: Display and analyze ROC curves. (2021).

43. Kembel, S. W., et al. picante: Integrating phylogenies and ecology. (2020).

67. Kembel, S. W., Wu, M., Eisen, J. A. & Green, J. L. Incorporating 16S gene copy number information improves estimates of microbial diversity and abundance. PLOS Comput. Biol. 8, e1002743 (2012).

68. Enwall, K., Throbäck, I. N., Stenberg, M., Söderström, M. & Hallin, S. Soil resources influence spatial patterns of denitrifying communities at scales compatible with land management. Appl. Environ. Microbiol. 76, 2243–2250 (2010).

50. Söderström M, Lindén B. Using precision agriculture data for planning field experiments e experiences from a research farm in Sweden. 2004. St. Petersburg, Rusia, p 161e168.

70. Barbera, P. et al. EPA-ng: Massively parallel evolutionary placement of genetic sequences. Syst. Biol. 68, 365–369 (2019).

52. Czech L, Barbera P, Stamatakis A. Genesis and Gappa: processing, analyzing and visualizing phylogenetic (placement) data. Bioinformatics 2020; 36: 3263–3265.

53. Dolédec S, Chessel D, ter Braak CJF, Champely S. Matching species traits to environmental variables: a new three-table ordination method. Environ Ecol Stat 1996; 3: 143–166.

31. Legendre, P., Galzin, R. & Harmelin-Vivien, M. L. Relating behavior to habitat: Solutions to the fourth-corner problem. Ecology 78, 547–562 (1997).

72. Dray, S. et al. ade4: Analysis of ecological data: exploratory and euclidean methods in environmental sciences. (2023).

56. ter Braak CJF, Cormont A, Dray S. Improved testing of species traits–environment relationships in the fourth-corner problem. Ecology 2012; 93: 1525–1526.

57. Eisenhofer R, Odriozola I, Alberdi A. Impact of microbial genome completeness on metagenomic functional inference. ISME Commun 2023; 3: 1–5.

58. Hink L, Gubry-Rangin C, Nicol GW, Prosser JI. The consequences of niche and physiological differentiation of archaeal and bacterial ammonia oxidisers for nitrous oxide emissions. ISME J 2018; 12: 1084–1093.

59. Alonso-Sáez L, Waller AS, Mende DR, Bakker K, Farnelid H, Yager PL, et al. Role for urea in nitrification by polar marine Archaea. Proc Natl Acad Sci 2012; 109: 17989– 17994.

60. Guénard G, von der Ohe PC, de Zwart D, Legendre P, Lek S. Using phylogenetic information to predict species tolerances to toxic chemicals. Ecol Appl 2011; 21: 3178– 3190.

61. Djemiel C, Maron P-A, Terrat S, Dequiedt S, Cottin A, Ranjard L. Inferring microbiota functions from taxonomic genes: a review. GigaScience 2022; 11: giab090.

37. Maurer, L. M., Yohannes, E., Bondurant, S. S., Radmacher, M. & Slonczewski, J. L. pH regulates genes for flagellar motility, catabolism, and oxidative stress in *Escherichia coli* K-12. J. Bacteriol. 187, 304–319 (2005).

63. Krulwich TA, Sachs G, Padan E. Molecular aspects of bacterial pH sensing and homeostasis. Nat Rev Microbiol 2011; 9: 330–343.

64. Abdo AI, Xu Y, Shi D, Li J, Li H, El-Sappah AH, et al. Nitrogen transformation genes and ammonia emission from soil under biochar and urease inhibitor application. Soil Tillage Res 2022; 223: 105491.

40. Wang, L. & Xiong, X. Long-term organic manure application alters urease activity and ureolytic microflora structure in agricultural soils. Agronomy 12, 3018 (2022).

66. Hazard C, Prosser JI, Nicol GW. Use and abuse of potential rates in soil microbiology. Soil Biol Biochem 2021; 157: 108242.

67. IPCC. IPCC Guidelines for national greenhouse gas inventories. in vol. 4 Chapter 11 (2006).

68. Swenson NG. Phylogenetic imputation of plant functional trait databases. Ecography 2014; 37: 105–110.

69. Zaneveld JRR, Thurber RLV. Hidden state prediction: a modification of classic ancestral state reconstruction algorithms helps unravel complex symbioses. Front Microbiol 2014; 5.

70. Debastiani VJ, Bastazini VAG, Pillar VD. Using phylogenetic information to impute missing functional trait values in ecological databases. Ecol Inform 2021; 63: 101315.

71. Guénard G. A phylogenetic modelling tutorial using Phylogenetic Eigenvector Maps (PEM) as implemented in R package MPSEM (0.4-1). 2022.

72. Stull GW, Pham KK, Soltis PS, Soltis DE. Deep reticulation: the long legacy of hybridization in vascular plant evolution. Plant J 2023; n/a.

73. Swenson NG, Weiser MD, Mao L, Araújo MB, Diniz-Filho JAF, Kollmann J, et al. Phylogeny and the prediction of tree functional diversity across novel continental settings. Glob Ecol Biogeogr 2017; 26: 553–562.

