## Supplementary material for "Predicting genomic traits in ammonia-oxidizing archaea using phylogenetic signals"

This document includes:

Supplementary text 1. An ammonium transporter gene found in Nitrosocaldales (*amt-NC*)

Supplementary Fig. 1. Phylogenetic tree based on amino acid alignment of the ammonium transporter gene (*amt*)

Supplementary Fig. 2. Segment of the Amt amino acid alignment

Supplementary Fig. 3. Updated reference phylogeny of the *amoA* gene

Supplementary Fig. 4. Association between strength of the phylogenetic signal (D) and accuracy of the predictions

Supplementary Fig. 5. Summary table of the fourth-corner analysis.

Supplementary Fig. 6. Association between genome and MAG completeness and validation parameters.

Supplementary Table 1. Correspondence between the *Nitrosocaldales* genomes and MAGs from Luo et al. [1] and the ones of this study.

Supplementary Table 2. Optimized parameters for the phylogenetic eigenvector-based predictive models of the genomic traits.

Supplementary Table 3. Accuracy, sensitivity and specificity of the predictions of genomic traits for each clade.

**Supplementary Text 1. An ammonium transporter gene found in Nitrosocaldales (*amt-NC*)**

When screening the AOA genomes and MAGs for the presence of the selected genes, we aligned each of the gene hits on the genomes and MAGs with the reference protein alignment from the EgNOGG database to double check that the hit was not an artefact. In the case of the Ammonium transporter genes (*amt*), aligning the hits with the reference protein alignment further allowed us to assign the hit to one of the two known ammonium transporter types [2–4].

When exploring the phylogenetic tree, and comparing it to the one reported by Offre et al.[4] we found an additional *amt* cluster in addition to the two groups reported in Fig.2 of Offre et al. [4]. This additional cluster (hereinafter *amt-NC*) was supported by a bootstrap value of 100/100, and contained 11 sequences, all of them belonging to genomes/MAGs of the Nitrosocaldales lineage (Supplementary Fig. 1). Among these sequences, there was *Nitrosocaldus islandicus* [5] and all the MAGs from the study of Luo et al. [1] All taxa having the *amt-NC* gene, also had the *amt2* gene (see Fig. 1a of main text).

The *amt-NC* ammonium transporter gene had two subgroups, hereinafter *amt-NC.1* and *amt-NC.2* which corresponded to the groups into which Luo et al. [1] divided the *Nitrosocaldales* lineage based on the concatenation of 122 archaeal genes. Specifically, GCA_013538795.1 and GCA_013538715.1, both having *amt-NC.1*, correspond to the *JZ-2.bins.172* and *QQ.bins.88* taxa, respectively, classified as clade A (see Fig. 1 from Luo et al.[1]). The rest of the genomes and MAGs that have *amt-NC.2* correspond to clades B, C, and D from Luo et al.[1] (Supplementary Table 1). It, therefore, seems that the two different *amt-NC* genes are related to the evolutionary history of the *Nitrosocaldus* lineage.

The *amt* amino acid alignment evidenced some differences between the two Amt-NC, and what Offree et al. [4] named Amt1 and Amt2 (Supplementary Fig. 2). The *amt-NC* contains the same amino acid as Amt1 and 2 at the ammonium-binding sites (arrows), the same histidine lining the transporter pore (triangle), and the same amino acids in many conserved loci (asterisks) [4]. However, *amt-NC* differs in several loci across the regions that Offre et al.[4] described in Figure 3 of their study (Supplementary Fig. 2). Altogether, these findings indicates that there exists a new ammonium transporter that may be unique for *Nitrosocaldales*. For this reason, we modeled the *amt-NC* separately in the phylogeny-based predictions. Future studies should examine in depth the functional differences between the proteins encoded by the *amt-NC, amt1* and *amt2* genes and compare, for example, their thermal stability.

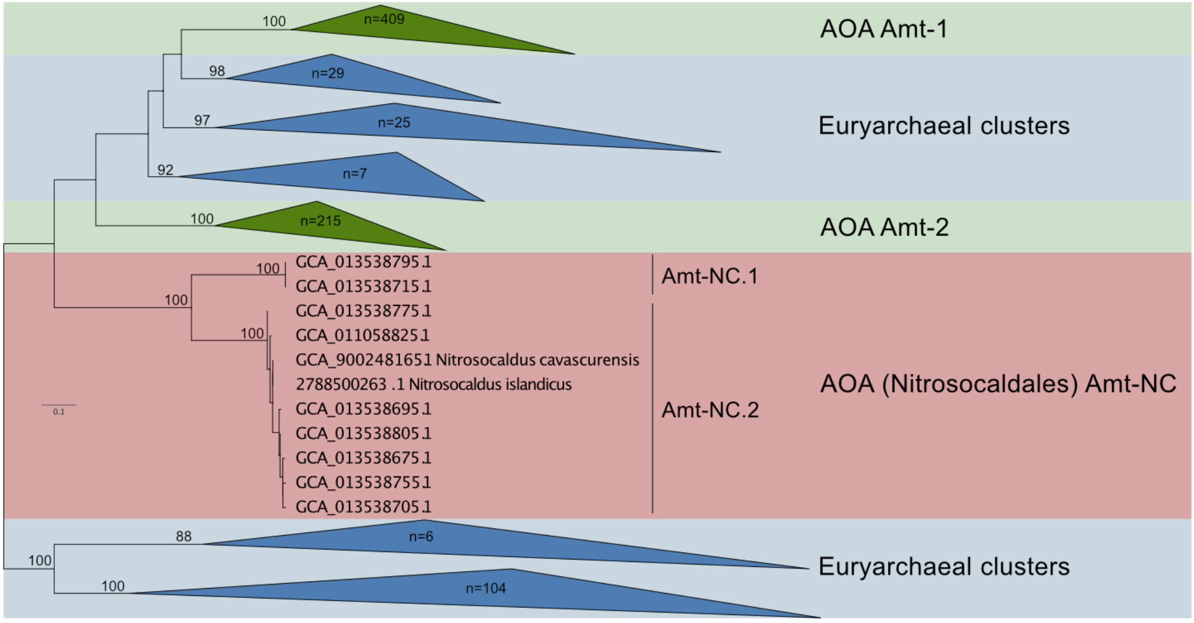


**Supplementary Fig. 1. Phylogenetic tree based on amino acid alignment of the ammonium transporter gene (*amt*).** The phylogenetic tree contains all genomes and MAGs of ammonia oxidizing archaea downloaded from NCBI and JGI, both with high and low completeness, as well as the sequences from the reference alignment of the arCOG04397 in the EggNOG database. It must be noted that the nomenclature for the ammonium transporters type 1 and 2 is the one from Offre et al. [4] .

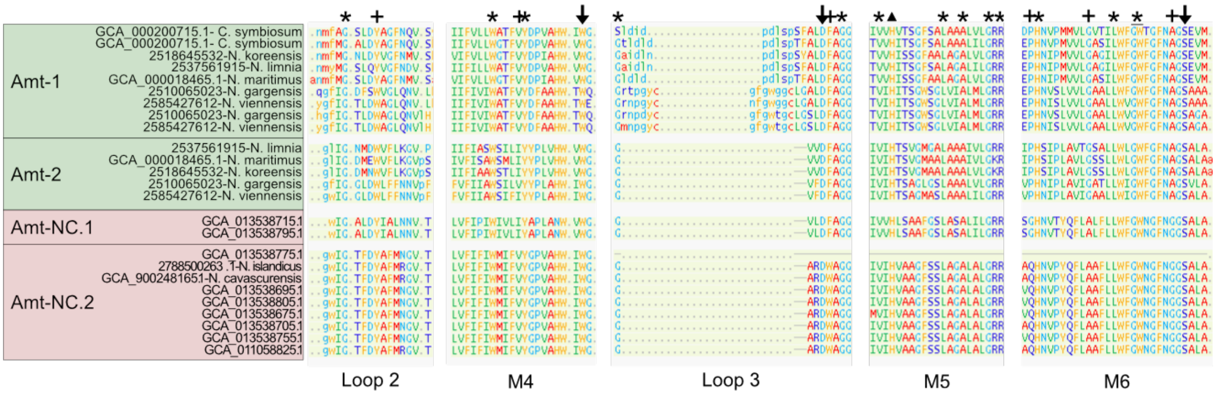


**Supplementary Fig. 2. Segment of the Amt amino acid alignment.** The figure contains the 11 sequences of Amt-NC and the Amt sequences from the taxa included in Offre et al.[4]. The regions are the ones described by Offre et al.[4]consisting of extracellular loops and transmembrane domains (M). Arrows depict the proposed ammonium-binding sites of Offre et al.[4]. Triangle depicts the first of the two conserved histidines lining the transporter pore of Offre et al.[4].Asterisks indicate loci with the same amino acid across the three *Amt* transporters, whereas crosses indicate loci with differences between them. It must be noted that the nomenclature for the ammonium transporters type 1 and 2 is the one from Offre et al.[4].

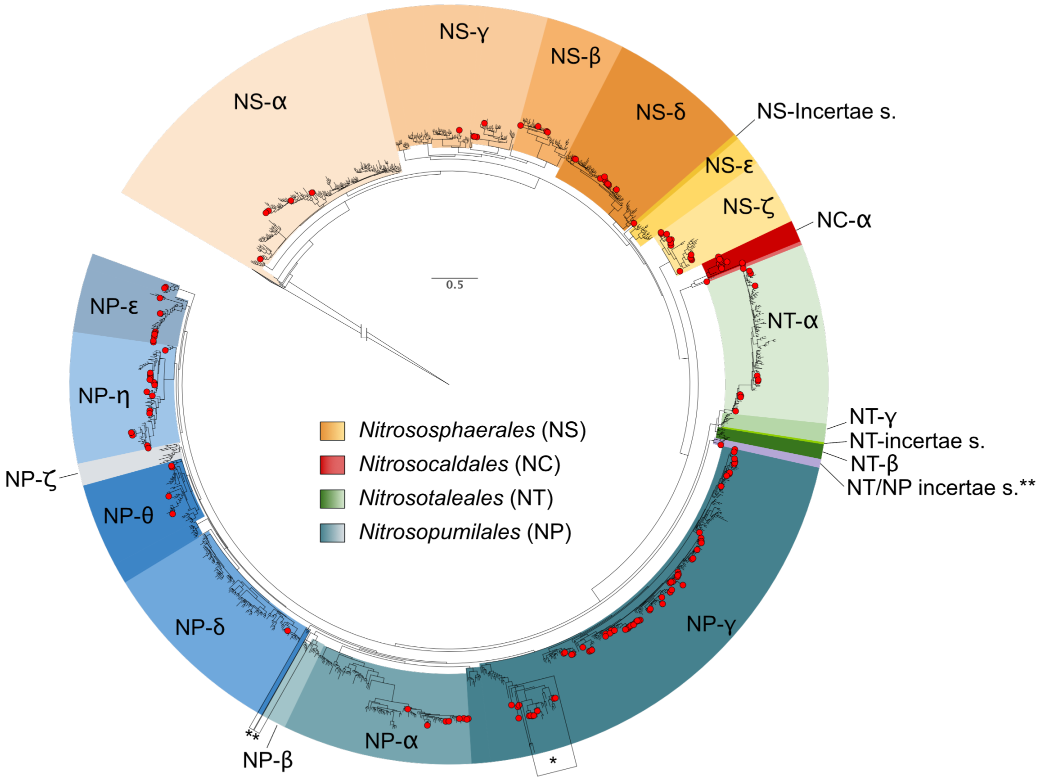


**Supplementary Fig. 3**. **Updated reference phylogeny of the *amoA* gene.** The phylogeny contains the archaeal *amoA* sequences from Alves et al. [6], the genomes and MAGs accessed at NCBI and JGI in the present study, and seven *amoA* sequences of ammonia-oxidizing bacteria (AOB) as an outgroup. Red circles are the available genomes and MAGs with > 80% completeness and < 5% contamination. A single asterisk (*) depicts the branches with differing topology between our updated reference phylogeny and that from Alves et al. [6]. Two asterisks (**) depict the *NT-NP incertae sedis* that is named *NP-ι* in Kerou et al. [7]. The tree is rooted on a collapsed gray clade of bacterial *amoA*.

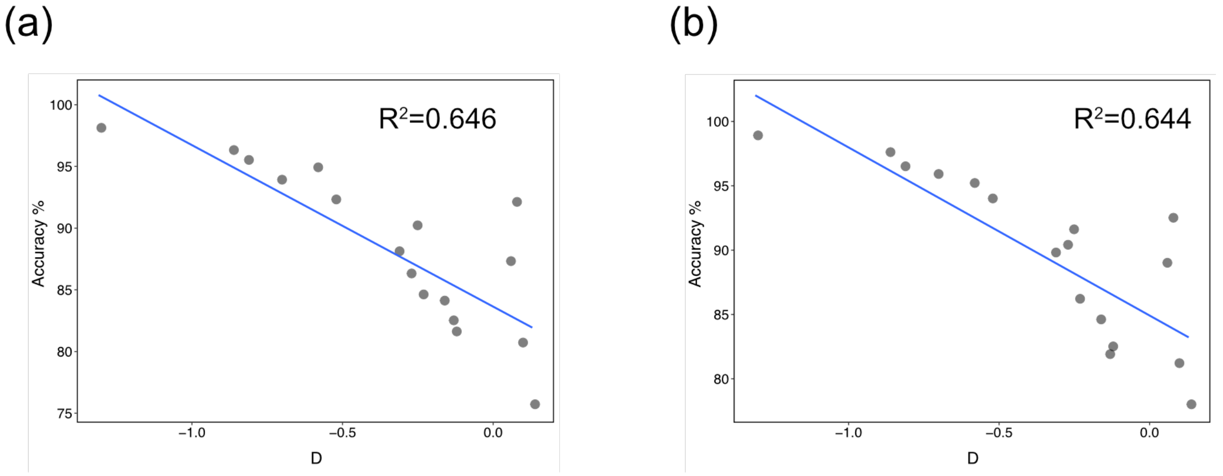


**Supplementary Fig. 4. Association between strength of the phylogenetic signal (D) and accuracy of the predictions**. **(a)** Phylogenetic eigenvector-based predictions and **(b)** ancestral state reconstruction.

(a)

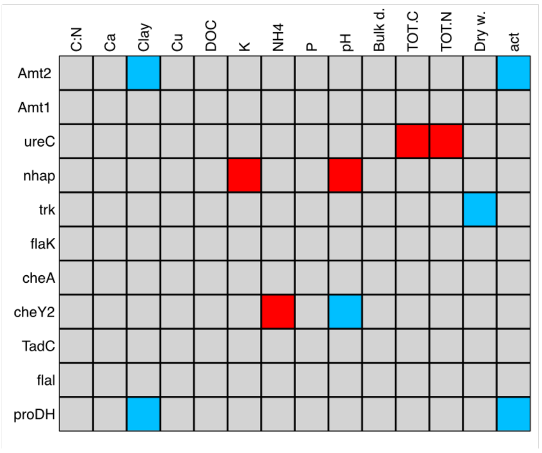


(b)

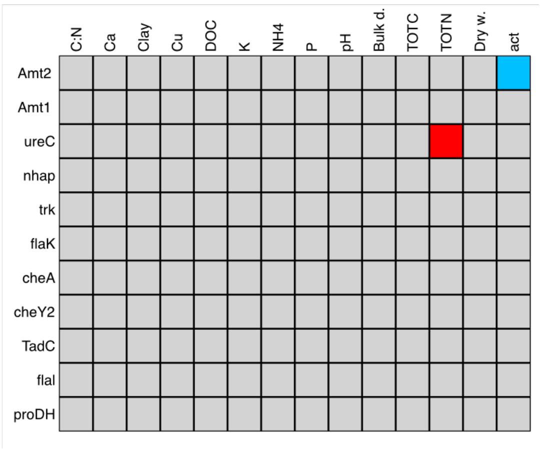


**Supplementary Fig. 5. Summary table of the fourth-corner analysis.** Grey squares represent non-statistically significant correlations, whereas red indicate positive correlations, and blue indicate negative correlations. **(a)** Correlations for non-corrected *P*-values ($P<0.05$) and **(b)** false discovery rate corrected *P*-values ($P < 0.065$).

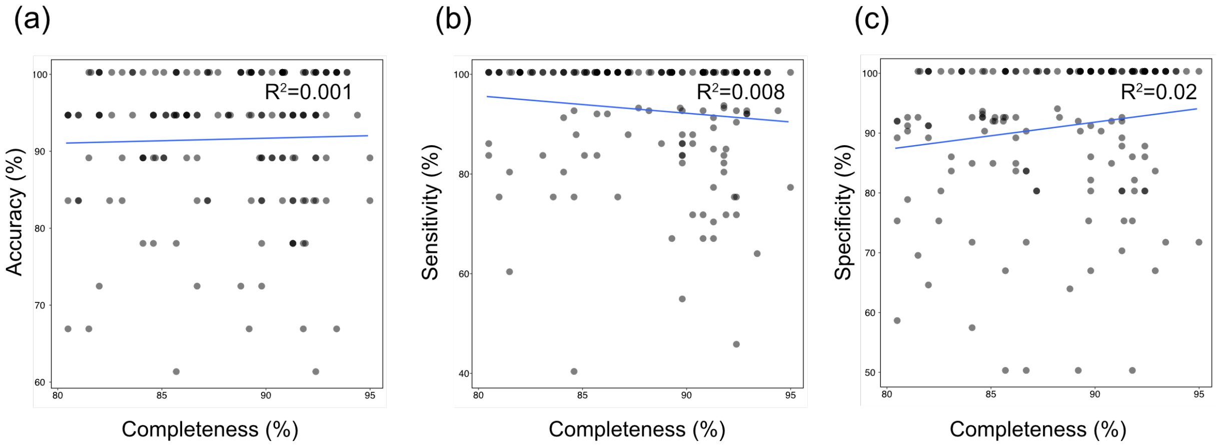


**Supplementary Fig. 6. Association between genome and MAG completeness and validation parameters**. Panels show prediction **(a)** accuracy, **(b)** sensitivity, and **(c)** specificity across the range of completeness of genomes and MAGs used in this study.

**Supplementary Table 1.** Correspondence between the *Nitrosocaldales* genomes and MAGs from Luo et al. [1] and the ones of this study.

| Genome_ID | Taxonomy | Name in Luo et al.[4] | Clade in Luo et al.[4] |
|---|---|---|---|
| GCA_013538795.1 | Ca. *Nitrosocaldus* sp. | JZ-2.bins.172 | A |
| GCA_013538715.1 | Ca. *Nitrosocaldus* sp. | QQ.bins.88 | A |
| GCA_013538775.1 | Ca. *Nitrosocaldus* sp. | JZ-2.bins.249 | D |
| GCA_011058825.1 | Ca. *Nitrosocaldus* sp. | Tumba.bins.72 | C |
| GCA_900248165.1 | Ca. *Nitrosocaldus cavascurensis* | *Nitrosocaldus cavascuriensis* | C |
| 2788500263 | Ca. *Nitrosocaldus islandicus* | *Nitrosocaldus islandicus* | C |
| GCA_013538695.1 | Ca. *Nitrosocaldus* sp. | JZ-3.bins.102 | D |
| GCA_013538805.1 | Ca. *Nitrosocaldus* sp. | JZ-1.bins.77 | D |
| GCA_013538675.1 | Ca. *Nitrosocaldus* sp. | QQ.bins.115137 | B |
| GCA_013538755.1 | Ca. *Nitrosocaldus* sp. | SRBZ.bins.174 | B |
| GCA_013538705.1 | Ca. *Nitrosocaldus* sp. | QQ.bins.97 | D |

**Supplementary Table 2. Optimized parameters for the phylogenetic eigenvector-based predictive models of the genomic traits.**

| Gene | $\lambda$* | Threshold** |
|---|---|---|
| *amt2* | 0.223511 | 0.859292 |
| *amt1* | 0.032208 | 0.442305 |
| *amt-NC* | 0.144659 | 0.122259 |
| *ureC* | 0.063092 | 0.682318 |
| *metE* | 0.059086 | 0.156239 |
| *metE2* | 0.064662 | 0.16398 |
| *proDH* | 0.044461 | 0.496194 |
| *rocA* | 0.11288 | 0.732061 |
| *cheA* | 0.030481 | 0.204702 |
| *cheY* | 0.06849 | 0.699686 |
| *cheY2* | 0.069469 | 0.589415 |
| *tadC* | 0.019416 | 0.798208 |
| *flaK* | 0.039111 | 0.655289 |
| *flaI* | 0.043561 | 0.543316 |
| *ipct* | 0.081492 | 0.157593 |
| *nhaP* | 0.132241 | 0.813169 |
| *trk* | 0.006893 | 0.775771 |
| *cspC* | 0.037884 | 0.234568 |

* Lambda ($\lambda$) is the tuning parameter for the shrinkage penalty of the elastic net regularisation.

** Threshold is the value above or below which probabilities were classified as presence or absence, respectively.

**Supplementary Table 3**. **Accuracy, sensitivity and specificity of the predictions of genomic traits for each clade.**

| Lineage | Clade | mean accuracy | mean sensitivity | mean specificity | total number of genomes/MAGs |
|---|---|---|---|---|---|
| NC | | 77.8% | 76.9% | 80.0% | 1 |
| NC | α (Alpha) | 97.2% | 99.2% | 94.3% | 10 |
| NP | α (Alpha) | 95.8% | 100.0% | 94.2% | 8 |
| NP | δ (Delta) | 77.8% | 80.0% | 75.0% | 1 |
| NP | ε (Epsilon) | 95.5% | 90.0% | 97.5% | 11 |
| NP | η (Eta) | 88.9% | 94.3% | 83.4% | 17 |
| NP | γ (Gamma) | 89.6% | 90.0% | 90.6% | 67 |
| NP | θ (Theta) | 90.3% | 92.9% | 82.5% | 4 |
| NS | α (Alpha) | 87.8% | 93.8% | 84.3% | 5 |
| NS | β (Beta) | 90.3% | 93.8% | 89.0% | 4 |
| NS | δ (Delta) | 93.2% | 93.7% | 93.5% | 9 |
| NS | γ (Gamma) | 96.7% | 100.0% | 95.3% | 5 |
| NS | *Incertae* | 88.9% | 100.0% | 84.6% | 1 |
| NS | ζ (Zeta) | 97.4% | 97.8% | 97.6% | 13 |
| NT | α (Alpha) | 91.4% | 93.7% | 91.2% | 11 |
| NT-NP | *Incertae* | 77.8% | 71.4% | 81.8% | 1 |

**Supplementary References**


5. Luo, Z.-H. *et al.* Genomic insights of "Candidatus Nitrosocaldaceae" based on nine new metagenome-assembled genomes, including "Candidatus Nitrosothermus" gen nov. and two new species of "Candidatus Nitrosocaldus". *Frontiers in Microbiology* **11**, (2021).

1. Nakagawa, T. & Stahl, D. A. Transcriptional response of the archaeal ammonia oxidizer *Nitrosopumilus maritimus* to low and environmentally relevant ammonia concentrations. *Applied and Environmental Microbiology* **79**, 6911–6916 (2013).

3. Wright CL, Lehtovirta-Morley LE. Nitrification and beyond: metabolic versatility of ammonia oxidising archaea. *ISME J* 2023; **17**: 1358–1368.

4. Offre P, Kerou M, Spang A, Schleper C. Variability of the transporter gene complement in ammonia-oxidizing archaea. *Trends Microbiol* 2014; **22**: 665–675.

4. Daebeler, A. *et al.* Cultivation and genomic analysis of 'Candidatus *Nitrosocaldus islandicus*,' an obligately thermophilic, ammonia-oxidizing Thaumarchaeon from a hot spring biofilm in Graendalur Valley, Iceland. *Front Microbiol* **9**, 193 (2018).

6. Alves RJE, Minh BQ, Urich T, Haeseler A von, Schleper C. Unifying the global phylogeny and environmental distribution of ammonia-oxidising archaea based on amoA genes. *Nat Commun* 2018; **9**: 1–17.

7. Kerou M, Ponce-Toledo RI, Zhao R, Abby SS, Hirai M, Nomaki H, et al. Genomes of Thaumarchaeota from deep sea sediments reveal specific adaptations of three independently evolved lineages. *ISME J* 2021; **15**: 2792–2808.
